# Ovarian Cancer Drives TLR5-Dependent Expansion of Myeloid Progenitors Through Systemic Dissemination of Ligands

**DOI:** 10.1101/2025.06.10.658497

**Authors:** Sree H. Kolli, Mitchell T. McGinty, Audrey M. Putelo, Cara N. Hatzinger, Simona Bajgai, Mika K. Poblete, Brandon Thompson, Tzu-Yu Feng, Francessca N. Azar, Akshita Mirani, William A. Petri, Stacey L. Burgess, Melanie R. Rutkowski

## Abstract

Ovarian cancer remains the most lethal gynecologic malignancy, due in part to the establishment of a profoundly immunosuppressive tumor microenvironment (TME). While TLR5 signaling has previously been implicated in promoting myeloid cell recruitment to the ovarian TME, the upstream source of ligand and its systemic effects on hematopoiesis remain poorly understood(1,2). Here, we show that ovarian cancer disrupts gut barrier integrity, leading to systemic translocation of TLR5 ligands into the peritoneum, blood, and bone marrow. This translocation correlates with enhanced expansion of myeloid progenitors in the bone marrow of wild-type (WT) but not TLR5-deficient (TLR5 KO) mice, leading to enhanced accumulation of monocytes into the tumor microenvironment. Pharmacologic blockade of TLR5 in tumor-bearing mice alters the composition of tumor-associated myeloid populations, increasing the frequency of monocytes and CCR2-expressing macrophages In the bone marrow of tumor-bearing WT mice. In the bone marrow, blockade of TLR5 signaling led to expansion of granulocyte-monocyte progenitors (GMPs), a phenotype recapitulated in a competitive chimera model. In vitro, stimulation of WT bone marrow cells with purified TLR5 ligands led to enhanced colony formation and skewed differentiation toward granulocyte-macrophage lineages. These data reveal that chronic TLR5 signaling, driven by tumor-induced gut leakage, promotes expansion of myeloid cells within the bone marrow and is a host-intrinsic mechanism driving accumulation of immature monocytes and macrophages into the tumor microenvironment.

## Introduction

Among women in the United States, ovarian cancer (OC) is the most lethal gynecologic malignancy with an estimated 19,880 new diagnosis and 12,740 deaths in 2024(3). In spite of surgical cytoreduction and platinum-based chemotherapy, ovarian cancer often spreads and relapses with poor prognosis(4). The tumor microenvironment (TME) plays a central role in ovarian cancer progression, driven by a complex and heterogeneous interplay of immune cells that vary across tumor stage and anatomical site. Some tumors are immunologically “deserted,” while others exhibit immune infiltration either peripherally or intratumorally(5). Macrophages and T lymphocytes—particularly CD3+, CD4+, and CD8+ T cells—are frequently observed in the ovarian TME, though their abundance and localization change with disease progression(6). Macrophages are often the most prevalent immune cell population in both tumor tissue and ascitic fluid and are known to promote immune suppression and tumor growth(7,8).The TME also includes regulatory T cells, myeloid-derived suppressor cells (MDSCs), and platelets, which can promote tumor growth and immune evasion(9). In cancer, the rise of pro-tumorigenic cell types and reduction of anti-tumorigenic cell types is influenced by emergency myelopoiesis.

Emergency myelopoiesis is a process of inflammation-induced hematopoiesis that replenishes myeloid cells during infection(10). It involves the expansion and differentiation of hematopoietic stem cells (HSCs) and myeloid progenitors, and is driven by inflammatory cytokines, interferons, and transcription factors(10,11). In cancer, this process is characterized by sustained production of inflammatory factors leading to the expansion of immature myelomonocytic and granulocytic myeloid cells from the bone marrow, that when they enter into the tumor microenvironment, are reprogrammed to support tumor growth and immune suppression(12). Tumor-derived growth factors—including granulocyte colony-stimulating factor (G-CSF), granulocyte-macrophage colony-stimulating factor (GM-CSF), and macrophage colony-stimulating factor (M-CSF/CSF1)—promote the expansion and lineage biasing of myeloid progenitors within the bone marrow(13). These progenitors give rise to immunosuppressive myeloid-derived suppressor cells (MDSCs), whose frequency is elevated in tumor-bearing hosts(12,13). Once expanded, MDSCs are recruited to the tumor microenvironment by chemokines such as CCL2 and CXCL1 and inflammatory cytokines like IL-6, where they are further conditioned by local signals— including hypoxia, TGF-β, and prostaglandins—to adopt a tumor-promoting phenotype(14–16). Thus, the immunosuppressive activity of myeloid cells in cancer arises through a multistep process that includes altered progenitor differentiation in the bone marrow, chemokine-mediated recruitment to the tumor, and functional polarization within the TME.

Recently we have shown that toll-like receptor 5 (TLR5) plays a complex role in ovarian cancer progression and immunotherapy response through reshaping of the ovarian TME(1,2). TLR5 is a key component of the innate immune system that recognizes bacterial flagellin(17). TLR5 signaling consists of activation of the MYD88 signaling cascade leading to the activation of NF-kB and MAPK pathways(18,19). In ovarian cancer, TLR5-dependent flagellated bacteria can drive malignant progression by increasing systemic IL-6, which mobilizes myeloid-derived suppressor cells (MDSCs), culminating in a loss of anti-tumor T cell function and subsequent failure of immune therapy response(1,2). However, the absence of TLR5 signaling enhances the efficacy of anti-PD-L1 therapy in orthotopic models of ovarian cancer, leading to sustained survival and protection against tumor rechallenge(1).

Here, we investigate the mechanistic link between TLR5 signaling and the emergence of immunosuppressive myeloid populations in the ovarian tumor microenvironment, focusing on the bone marrow as a critical site of early myeloid programming. Previous studies have shown that TLR5 signaling influences myeloid populations within the ovarian TME(1,2). However, it remains unclear how ovarian cancer alters the availability of TLR5 ligands, and how these ligands, once disseminated systemically, may act on hematopoietic stem and progenitor cells (HSPCs) to contribute to emergency myelopoiesis. We address this gap by examining how tumor-associated increases in gut permeability—observed in both WT and TLR5-deficient mice—permit translocation of TLR5 ligands to distal compartments, including the bone marrow. We find that exposure to flagellin is associated with expansion of granulocyte-monocyte progenitors (GMPs), and that this expansion is dependent on TLR5 expression. Using a competitive bone marrow chimera model, we demonstrate that in tumor-bearing hosts, there is an expansion of wild type, but not TLR5-deficient, GMP and macrophage/monocytic precursors within the bone marrow, culminating in increased numbers of monocytes within the tumor microenvironment. *In vitro* methylcellulose assays demonstrate that TLR5 stimulation enhances colony formation of granulocyte-macrophage lineages, supporting a functional role for TLR5 in the expansion of myeloid progenitors. Together, these findings suggest that TLR5 signaling, through tumor-associated gut leakage, contributes to the development of an immunosuppressive myeloid compartment by acting at the level of bone marrow progenitor differentiation.

## Materials & Methods

### Mice

TLR5 wild-type mice were generated using transgenic *Kras^tm4Tyj^*and *Trp53^tm1Brn^* mice(20,21) obtained from the National Cancer Institute (NCI) Mouse Models of Human Cancers Consortium and brought to a full C57BL/6 background(22). These mice were then bred to TLR5-deficient (TLR5 KO) mice (B6.129S1-*Tlr5^tm1Flv^*/J), as previously described(23), to generate TLR5 KO mice. CD45.1 mice (B6.SJL-*Ptprc^a^Pepc^b^*/BoyCrl) (24) were obtained from Charles River. All experiments were conducted utilizing adult (∼20-week-old) female mice. All strains were maintained in specific-pathogen-free barrier facilities at the University of Virginia. All experiments in this study were approved by the University of Virginia Institutional Animal Care and Use Committee.

### In Vivo TLR5 Inhibition

For *In vivo* inhibition of TLR5, 100µg/mouse of TH1020 small molecule inhibitor was injected IP for four consecutive days starting 10 days post-tumor initiation(25).

### Cell lines and implantation

ID8 cells were provided by K. Roby (Department of Anatomy and Cell Biology, University of Kansas) and retrovirally transduced to express *Defb29* and *Vegf-A*(*26*). ID8-*Defb29/Vegf-A* is an aggressive ovarian tumor cell line that recapitulates stage III/IV ovarian cancer.

Cell lines were authenticated by monitoring of morphology and monthly testing for mycoplasma. To limit the opportunity for genetic drift, cells were maintained at less than five passage numbers and maintained as frozen stocks at −180°C and expanded only for inoculation into mice. Tumor cell lines were cultured in RPMI complete media (RPMIc): RPMI (11875093, Gibco), 10% FBS (Sigma), 2 mmol/L of l-glutamine (25030081, Gibco), 1 mmol/L of sodium pyruvate (11360070, Gibco), 50 μmol/L of β-mercaptoethanol (M6250, Sigma), and 100 U/mL of Penicillin/Streptomycin (15140122, Gibco). ID8*-Defb29/Vegf-A* were initiated by intraperitoneal injection (IP) of 2e6 cells in sterile PBS at 100µl total volume.

### Gut Permeability Assay

Gut barrier integrity was assessed using a fluorescein isothiocyanate (FITC)-dextran assay(27). Mice received 4% dextran sodium sulfate (DSS) in drinking water ad libitum for 5 consecutive days prior to takedown(28). On day of assay mice were fasted for 4 hours prior to euthanasia, with continued access to water. FITC-dextran (4 kDa, Sigma-Aldrich) was prepared at 80 mg/mL in sterile 1× PBS and protected from light. Each mouse received 200 μL of the FITC-dextran solution by oral gavage using a sterile feeding needle. Exactly 1 hour after gavage, mice were euthanized by CO₂ asphyxiation followed by cervical dislocation. Blood was collected via cardiac puncture. Fluorescence intensity of the plasma was measured 96-well plates using a fluorescence plate reader (excitation: 485 nm; emission: 530 nm).

### Mouse TLR5 Reporter HEK293 reporter assay

Quantification of TLR5 ligands was performed using HEK-Blue™ mTLR5 cells (InvivoGen), which express murine TLR5 and secrete secreted embryonic alkaline phosphatase (SEAP) upon receptor activation(29). SEAP levels were measured using HEK-Blue™ Detection media according to the manufacturer’s instructions(29). A standard curve was generated using recombinant, ultra-purified *Salmonella typhimurium* flagellin (RecFLA-ST; InvivoGen). Serial dilutions ranging from 0 to 2000 pg/mL were prepared in endotoxin-free water and plated at 20 μL per well in a flat-bottom 96-well plate. Negative controls included endotoxin-free water and HEK-Blue™ Detection media alone. Peritoneal fluid was collected by flushing the peritoneal cavity with 5 mL of sterile PBS, followed by centrifugation at 500 × g for 5 minutes at 4°C to remove cells and debris. Bone marrow supernatants were prepared by flushing femurs with sterile PBS, followed by a 5-minute centrifugation at 500 × g to pellet cells.

HEK-Blue™ mTLR5 cells were washed with pre-warmed PBS and gently detached without enzymatic treatment. Cells were resuspended in HEK-Blue™ Detection media at a concentration of 140,000 cells/mL, and 180 μL (∼25,000 cells) was added to each well containing standards, controls, or 20 μL of processed sample. Plates were incubated at 37°C with 5% CO₂ for 16 hours. SEAP activity was measured by reading absorbance at 655 nm using a microplate spectrophotometer. Absorbance values from the standard curve were used to interpolate TLR5 ligand concentrations (pg/mL) in unknown samples.

### Flow Cytometry

Isolated tissues were placed on ice in a sterile 6-well plate with 3 mL of RMPI (11875093, Gibco) with 5% FBS (Sigma). Ascites were harvested via PBS wash of the peritoneal cavity by syringe aspiration followed by residual fluid collection by pipetting. To make single-cell suspensions before staining with antibodies, the digested tissues were passed through 70μm cell strainers (352350, Corning) using mechanical force with the rubber end of a 5mL syringe. For *in vitro* coculture experiments, all tissues were processed in sterile conditions.

For intracellular cytokine staining, disassociated tumor specimens were stained with the LIVE/DEAD fixable Aqua Dead Cell Stain Kit (Life Technologies). Cells were then fixed with 1% methanol formaldehyde solution (Thermo Scientific) followed by permeabilization in 0.5% Saponin solution (Sigma) and intracellular staining. Proliferation, surface, and intracellular staining were analyzed using FlowJo software. SPHERO™ AccuCount Particles (cat #ACFP-50-5) were utilized to enumerate cell counts. Flow cytometry experiments were performed on a Cytek Aurora Borealis (5 lasers) or Life Technologies Attune NxT.

Flow Cytometry data was analyzed using FlowJo version 10.10.0. Principal component analysis was performed using FlowJo-integrated UMAP analysis software. FlowSOM version 3.0.18 was downloaded from FlowJo exchange (https://www.flowjo.com/exchange/#/) and utilized to visualize and cluster high-parameter flow cytometry data using FlowJo.

### Mixed Bone Marrow Chimera

Bone marrow cells were prepared from femurs and tibias of TLR5 KO (CD45.2) and wild-type (CD45.1) donor mice. Recipient wild-type (CD45.1) mice were irradiated (2 consecutive days x 600 rads/day) and retro-orbitally injected with an equal 1:1 mix of donor bone marrow cells (TLR5 KO:wild-type). Tumors were initiated 10-weeks post-bone marrow reconstitution via IP injection of 2e6 ID8-*Defb29/Vegf-A* cells. Tissue samples were collected 15 days after tumor initiation and analyzed by flow cytometry.

### Colony Forming Assay

To assess hematopoietic colony formation, whole bone marrow was harvested from wild-type (WT) and TLR5 knockout (TLR5 KO) mice. Cells were resuspended in Iscove’s Modified Dulbecco’s Medium (IMDM) containing 2% fetal bovine serum (FBS) and plated into MethoCult™ GF M3434 methylcellulose-based medium (StemCell Technologies), which contains stem cell factor, IL-3, IL-6, and erythropoietin, according to the manufacturer’s protocol. A total of 20,000 unfractionated bone marrow cells were seeded per well in 6-well plates in triplicate under three conditions: unstimulated (media only), stimulated with ultra-purified flagellin from *S. typhimurium* (FLA-ST, 100 ng/mL, InvivoGen), or stimulated with lipopolysaccharide (LPS, 100 ng/mL, InvivoGen). Cultures were incubated at 37°C in 5% CO₂ for 11 days. On day 11, colonies were quantified manually under a light microscope. Colonies were classified into total colonies, burst-forming unit erythroid (BFU-E), granulocyte-erythrocyte-monocyte-megakaryocyte (GEMM), and granulocyte-macrophage (GM) based on morphological criteria. Scoring was performed with blinding to genotype and treatment condition.

## Results

### Ovarian Cancer and DSS Treatment Induce Gut Permeability and Translocation of TLR5 Ligands

Increased gut permeability has been reported in various cancers, including colorectal cancer, melanoma, orthotopic renal cancer, and fibrosarcoma, where tumor-induced barrier dysfunction leads to microbial translocation, dysbiosis, and systemic inflammation(30,31). Given these observations, we sought to determine whether ovarian cancer similarly disrupts gut barrier integrity and promotes the translocation of TLR5 ligands into circulation and distal tissues.

To assess gut permeability, we performed a FITC-Dextran assay(27) in wild-type (WT) and TLR5 knockout (TLR5 KO) mice with or without ID8-*Defb29/Vegf-A* tumors implanted orthotopically into the peritoneal environment for 20 days. Mice treated with 4% dextran sodium sulfate (DSS) in drinking water served as a positive control for chemically induced gut barrier disruption(28). On the day of analysis, mice were fasted for 4 hours and gavaged with 200 µL of 80 mg/mL 4 kDa FITC-dextran exactly 1 hour prior to euthanasia; blood was collected by cardiac puncture and fluorescence intensity in plasma was measured to quantify FITC-dextran translocation as an indicator of barrier permeability (**Figure 1A**). In the absence of tumor burden or DSS exposure, both WT and TLR5 KO mice exhibited low fluorescence intensity in peripheral blood, indicative of intact gut barrier function. However, in both tumor-bearing and DSS-treated mice, fluorescence intensity was significantly elevated, demonstrating increased gut permeability (**Figure 1B**). Importantly, there were no significant differences in permeability between WT and TLR5 KO mice under tumor or DSS conditions, suggesting that gut barrier disruption occurs independently of TLR5 signaling (**Figure 1B**).

**Figure 1:**
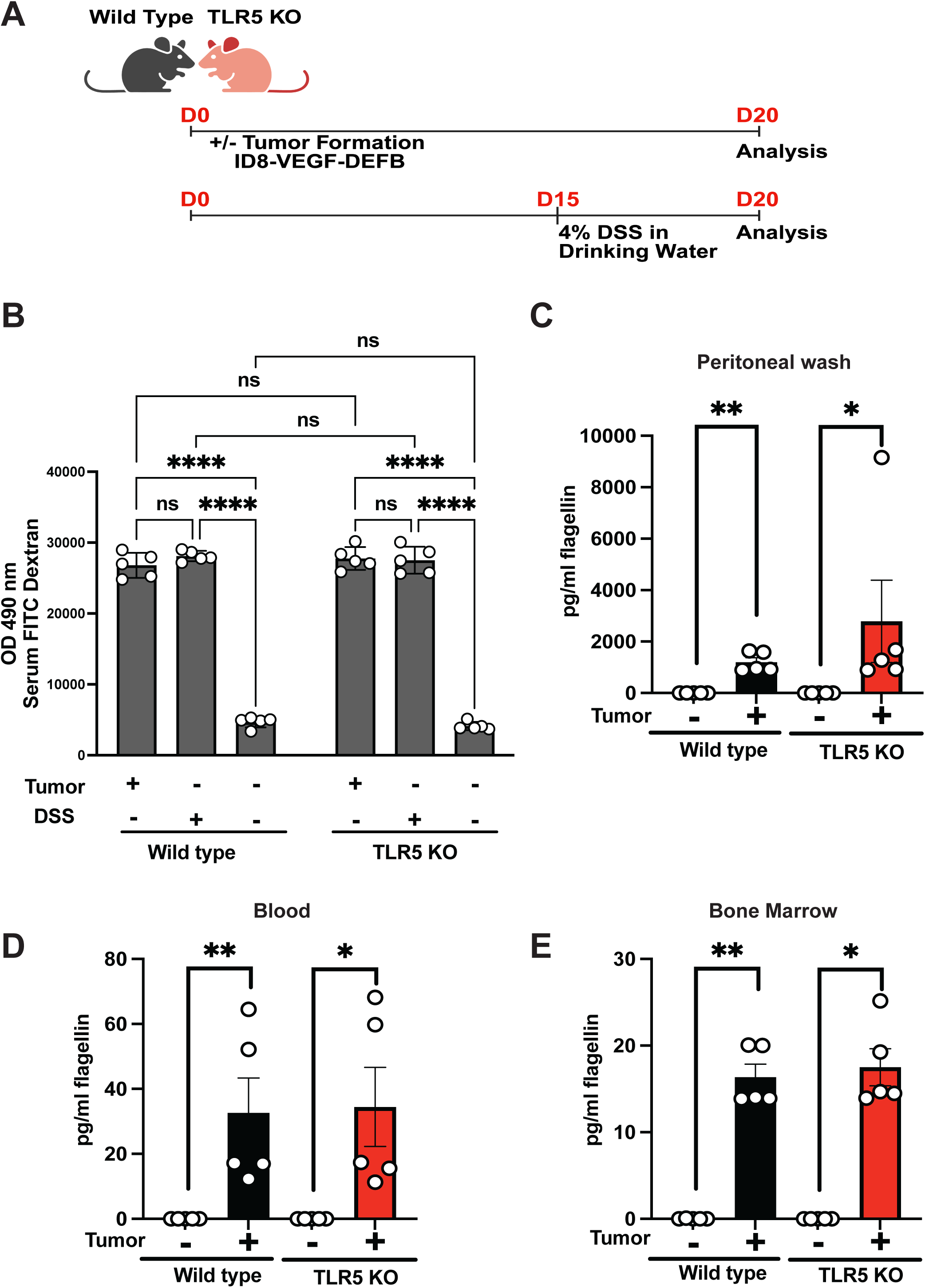
Ovarian cancer and DSS treatment induce gut permeability and translocation of TLR5 ligands. **(A)** Gut permeability was evaluated using a FITC-dextran assay in wild-type (WT) and TLR5 knockout (TLR5 KO) mice either left untreated, treated with dextran sodium sulfate (DSS, 4% in drinking water for 5 days), or implanted intraperitoneally with 2 × 10⁶ ID8-Defb29/VEGF-A ovarian tumor cells. Mice were fasted for 4 hours prior to oral gavage with 200 µL of 80 mg/mL FITC-dextran. One hour later, blood was collected via cardiac puncture, and fluorescence intensity was measured in plasma as an indicator of gut barrier disruption. **(B–D)** Quantification of translocated TLR5 ligands in WT and TLR5 KO mice using a HEK-Blue™ mTLR5 reporter cell line. Peritoneal fluid **(B),** peripheral blood **(C),** and bone marrow supernatant **(D)** were harvested from untreated and tumor-bearing mice at day 20 post tumor implantation. Reporter cells were stimulated with collected samples in HEK-Blue™ Detection medium, and SEAP activity was measured via absorbance at 655 nm following 16 hours of incubation. Flagellin concentrations (pg/mL) were calculated using a standard curve generated from a serial dilution (0–2000 pg/mL) of ultrapure recombinant flagellin. In all three compartments, tumor-bearing mice had significantly elevated levels of TLR5 ligands relative to their non-tumor counterparts, independent of TLR5 expression. Data represent individual mice with group means ± SEM. Statistical significance was determined using unpaired two-tailed t-tests; *p < 0.05, **p < 0.01.

To determine whether increased permeability facilitated the dissemination of bacterial ligands(32) and specifically TLR5 ligands, we quantified their presence in peritoneal fluid, blood, and bone marrow using a TLR5 reporter cell line(29) (**Figure 1C–E**). In mice bearing ID8-*Defb29/Vegf-A* tumors for 20 days we quantitated an increase in TLR5 ligands within the peritoneal wash/tumor microenvironment (**Figure 1C**), blood (**Figure 1D**), and bone marrow (**Figure 1E**), compared to non-tumor-bearing control animals. This pattern was observed in both WT and TLR5 KO mice, indicating that tumor-induced translocation of TLR5 ligands is not dependent on host TLR5 expression.

These findings establish that ovarian cancer, similar to other extraintestinal malignancies(30), compromises gut integrity and permits the systemic spread of microbial components, including TLR5 ligands. The presence of TLR5 ligands in distal tissues raises the possibility that microbial translocation may contribute to immune modulation in the tumor microenvironment, warranting further investigation into its potential role in shaping ovarian cancer progression.

### TLR5 signaling promotes accumulation of CCR2+ Ly6C+ myeloid cells in the ovarian tumor microenvironment

To investigate the role of TLR5 signaling in shaping the ovarian tumor myeloid landscape, we analyzed immune cell populations in mice bearing ID8-*Defb29/Vegf-A* tumors treated with TH1020, a specific small molecule inhibitor targeting TLR5(33), or vehicle control (**Figure 2A**). Treatment was initiated on day 10 post-tumor implantation and continued daily for four consecutive days prior to analysis on day 20. At the time of analysis, ascites fluid was collected and used to assess tumor-infiltrating immune cell composition with flow cytometry. Using UMAP on pre-gated CD11b+myeloid cells (**Supplemental Figure 1A**), distinct shifts in myeloid composition following TLR5 blockade were observed (**Figure 2B–C**). Cluster-level analysis and marker heatmaps indicated that the most prominent reduction occurred in CCR2+ Ly6C+ populations (**cluster 9; Figure 2D**), a subset of inflammatory monocytes known to be recruited from the bone marrow to sites of inflammation in a CCR2-dependent manner(34), and that associate with immune suppression and poor outcomes(35,36).

**Figure 2:**
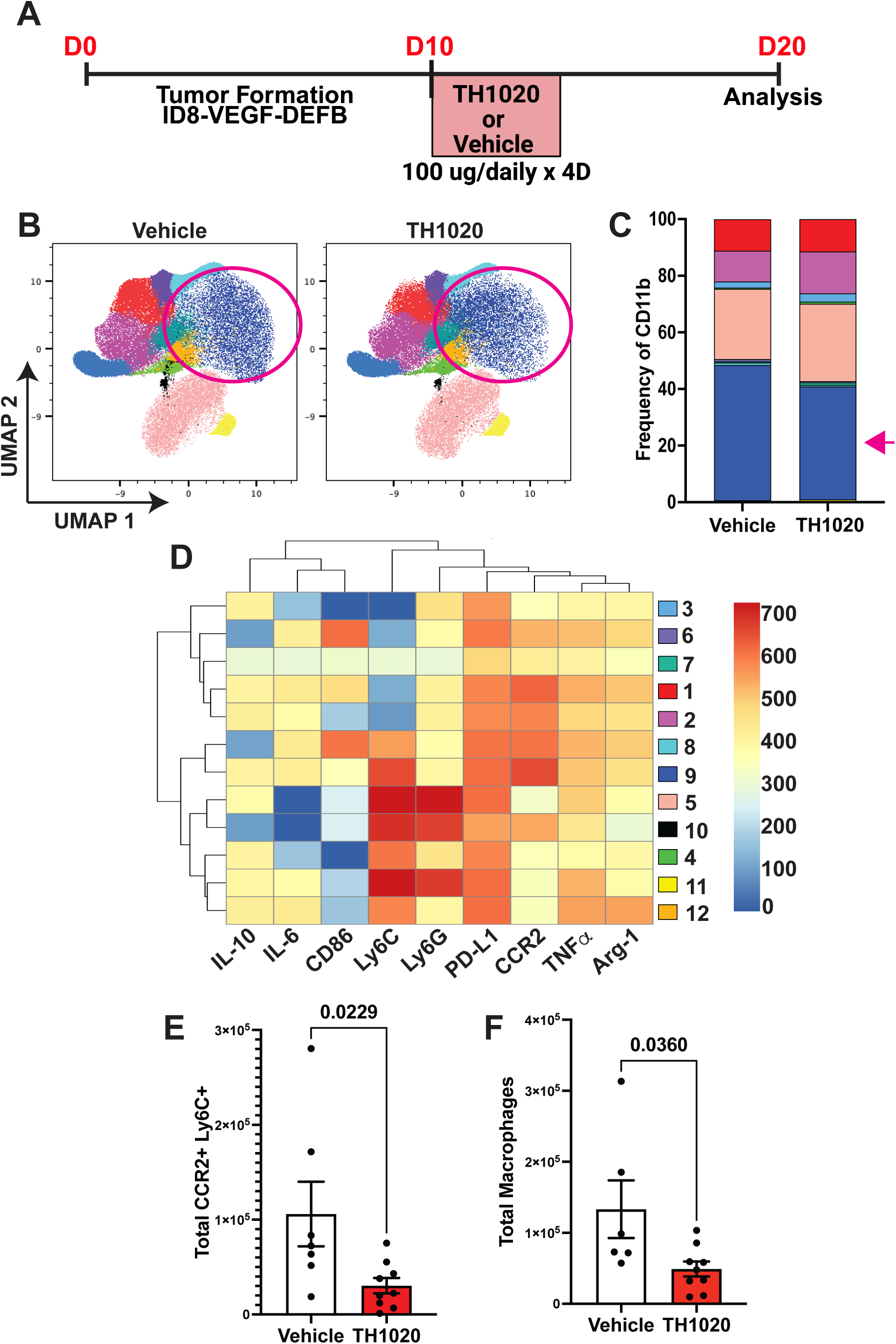
TLR5 blockade remodels the tumor-associated myeloid compartment and reduces the abundance of CCR2⁺Ly6C⁺ cells in the ovarian TME. **A)** UMAP projections of tumor-infiltrating CD11b⁺ myeloid cells isolated from ID8-Defb29/VEGF-A tumor-bearing mice treated with either vehicle or TLR5-blocking antibody (TH1020). Cells were pooled from n=5 mice per group and analyzed via high-parameter spectral flow cytometry. Annotated clusters were defined by unsupervised FlowSOM clustering and overlaid on UMAP projections; colored ovals highlight a dominant population reduced upon TLR5 blockade. **(B)** Cluster frequencies of CD11b⁺ myeloid cells from both treatment groups displayed as stacked bar graphs. Pink arrow denotes a marked decrease in Cluster 9 in TH1020-treated mice compared to vehicle controls. **(C)** Heatmap of median expression values for canonical myeloid markers across all clusters. Clusters are hierarchically ordered based on phenotypic similarity. Cluster 9 (highlighted with arrow) is characterized by high expression of Ly6C, CCR2, PD-L1, and Arg-1, consistent with monocytic myeloid-derived suppressor cells (M-MDSCs). **(D)** Absolute quantification of CCR2⁺Ly6C⁺ cells in tumor digests from vehicle and TH1020-treated mice. Cells were counted using fluorescent counting beads and gating on live, singlet, CD45⁺CD11b⁺Ly6C⁺CCR2⁺ populations. TLR5 blockade significantly reduced the abundance of CCR2⁺Ly6C⁺ cells in the TME (unpaired two-tailed t-test, p = 0.0229). Data shown as mean ± SEM with individual mouse values overlaid.

Validating the UMAP results using hierarchical gating (**supplemental Figure 1A**), we confirmed that CCR2+ Ly6C+ cells were significantly reduced within the TME after TH1020-treatment (**Figure 2E**). We also observed a decrease in total tumor-infiltrating macrophages (**Figure 2F**). Upon entering the TME, Ly6C+ CCR2+ monocytes can become tumor-associated macrophages with pro-tumorigenic functions(37,38), suggesting that blockade of TLR5 signaling affected a monocyte/macrophage axis within the ovarian TME. Supporting this rational, TLR5 blockade led to a significant reduction in CCR2+ Ly6C+ F4/80+ macrophages (**Supplemental Figure 1B–C**). Altogether, these data suggest that TLR5 signaling contributes to the accumulation of inflammatory monocytes and macrophages within the ovarian TME and indicate that TLR5 signaling influences the recruitment and local composition of myeloid cells in the ovarian TME.

### TLR5 Signaling Drives Tumor-Associated Myeloid Expansion through Bone Marrow Progenitor Skewing

Given that CCR2+ Ly6C+ cells are derived from the bone marrow, our data suggest that TLR5-mediated enrichment of myeloid cells in the TME, particularly tumor-associated macrophages, could be the result of enhanced output from bone marrow progenitor compartments during tumor progression(39,40). This phenotype may therefore reflect TLR5-mediated effects on myelopoiesis or progenitor expansion prior to tumor infiltration. Because we measured an increased presence of TLR5 ligands in the bone marrow of tumor-bearing animals, we next investigated whether TLR5 signaling contributes to early myeloid skewing in hematopoietic progenitors within the bone marrow. We hypothesized that TLR5 signaling facilitates this process by promoting expansion and differentiation of monocyte progenitors in the bone marrow.

To test this, we quantified progenitor populations in the femurs of wild-type (WT) with or without treatment using TH1020 and compared differences to TLR5 knockout (TLR5 KO) mice with or without ID8-*Defb29/Vegf-A* tumors (**Figure 3A**). Using the gating strategy outlined in **Supplemental 2A**, we identified that tumor-bearing WT mice had a significant expansion of common monocyte progenitors (CMoPs) compared to non-tumor-bearing controls, an effect that was abolished in both tumor-bearing TLR5 KO and TH1020-treated animals (**Figure 3B**). This suggests a TLR5-dependent expansion of monocyte progenitors during tumor progression.

**Figure 3:**
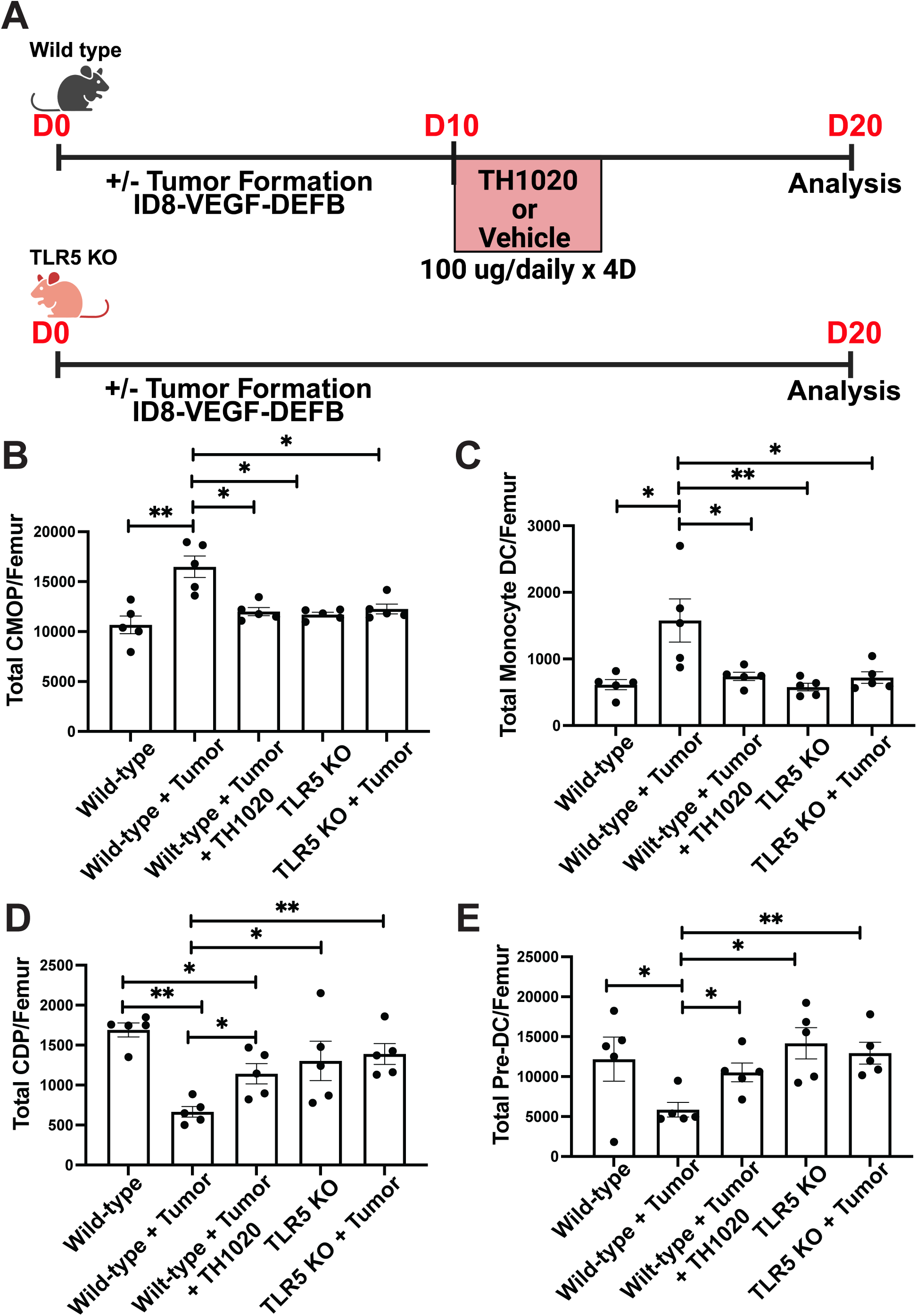
TLR5 signaling influences myeloid progenitor populations in the bone marrow of tumor-bearing mice. **(A)** Experimental schema depicting timelines for wild-type (WT) and TLR5 knockout (TLR5 KO) mice with or without orthotopic ovarian tumor implantation. Tumors were initiated intraperitoneally (i.p.) on day 0, and femurs were harvested on day 20 for flow cytometric analysis of hematopoietic progenitor populations. Five mice were used per group. **(B–F)** Quantification of total bone marrow cell counts per femur for key hematopoietic and myeloid progenitor subsets using multiparameter flow cytometry. Data shown for WT and TLR5 KO mice, with and without tumor burden. **(B)** Hematopoietic stem cells (HSCs) (Flow Gating Strategy) show no significant changes across genotypes or tumor status. **(C)** Common monocyte progenitors (CMPs) (Flow Gating Strategy) are significantly expanded in WT tumor-bearing mice relative to WT non-tumor controls, but this expansion is abrogated in TLR5 KO tumor-bearing mice. **(D)** Monocyte lineage dendritic cells (moDCs) (Flow Gating Strateg) are also significantly elevated in WT tumor-bearing mice compared to all other groups. **(E)** Common dendritic cell precursors (CDPs) (Flow Gating Strategy) are reduced in tumor-bearing WT mice, an effect not observed in TLR5 KOs. **(F)** Pre-classical dendritic cells (pre-cDCs) (Flow Gating Strategy) similarly decrease in tumor-bearing WT mice but remain unchanged in TLR5 KO mice. Bars represent mean ± SEM. Individual mouse data points shown. Statistical significance determined by two-way ANOVA with Tukey’s multiple comparison test. *p < 0.05, **p < 0.01.

Similarly, tumor-bearing WT mice exhibited elevated frequencies of monocyte-derived dendritic cells (moDCs), while exhibiting a marked reduction in common dendritic cell precursors (CDPs) and pre-classical dendritic cells (pre-cDCs), which upon TH1020 treatment were restored to levels similar to that observed in TLR5 KO tumor-bearing animals (**Figure 3C–E**). These changes occurred without any alteration in total hematopoietic stem cell (HSC) numbers (**Supplemental Figure 2B**), suggesting that TLR5 signaling influences the lineage commitment of myeloid precursors rather than maintenance of stem cells. Together, these findings support a model in which chronic TLR5 signaling in tumor-bearing hosts promotes expansion of bone marrow monocyte progenitors, seeding the TME with tumor-associated macrophages and myeloid suppressor populations.

### TLR5 Signaling Directly Regulates Myeloid Progenitor Differentiation in the Bone Marrow

Our previous findings suggested that TLR5 signaling promotes the accumulation of tumor-associated macrophages in the ovarian TME through effects on bone marrow progenitor populations. The possibility existed that differences occurring within the bone marrow were occurring in response to microenvironmental differences within the bone marrow. To determine whether TLR5 signaling directly regulates myeloid differentiation in the bone marrow, we generated competitive bone marrow chimeras by transplanting lethally irradiated CD45.1+ WT recipient mice with a 1:1 mixture of bone marrow from WT (CD45.1+) and TLR5 knockout (CD45.2+) donors. After 10 weeks of engraftment, mice were either left untreated or orthotopically implanted with ID8-Defb29/VEGF-A tumors, and the bone marrow, spleen and tumor microenvironment were analyzed 15 days later (**Figure 4A**). The gating strategy to evaluate the bone marrow is depicted in **Supplemental Figure 3A**. Engraftment efficiency and systemic chimerism were confirmed prior to tumor implantation, showing no significant differences between WT and TLR5 KO leukocyte compartments (**Supplemental Figure 3B**).

**Figure 4.**
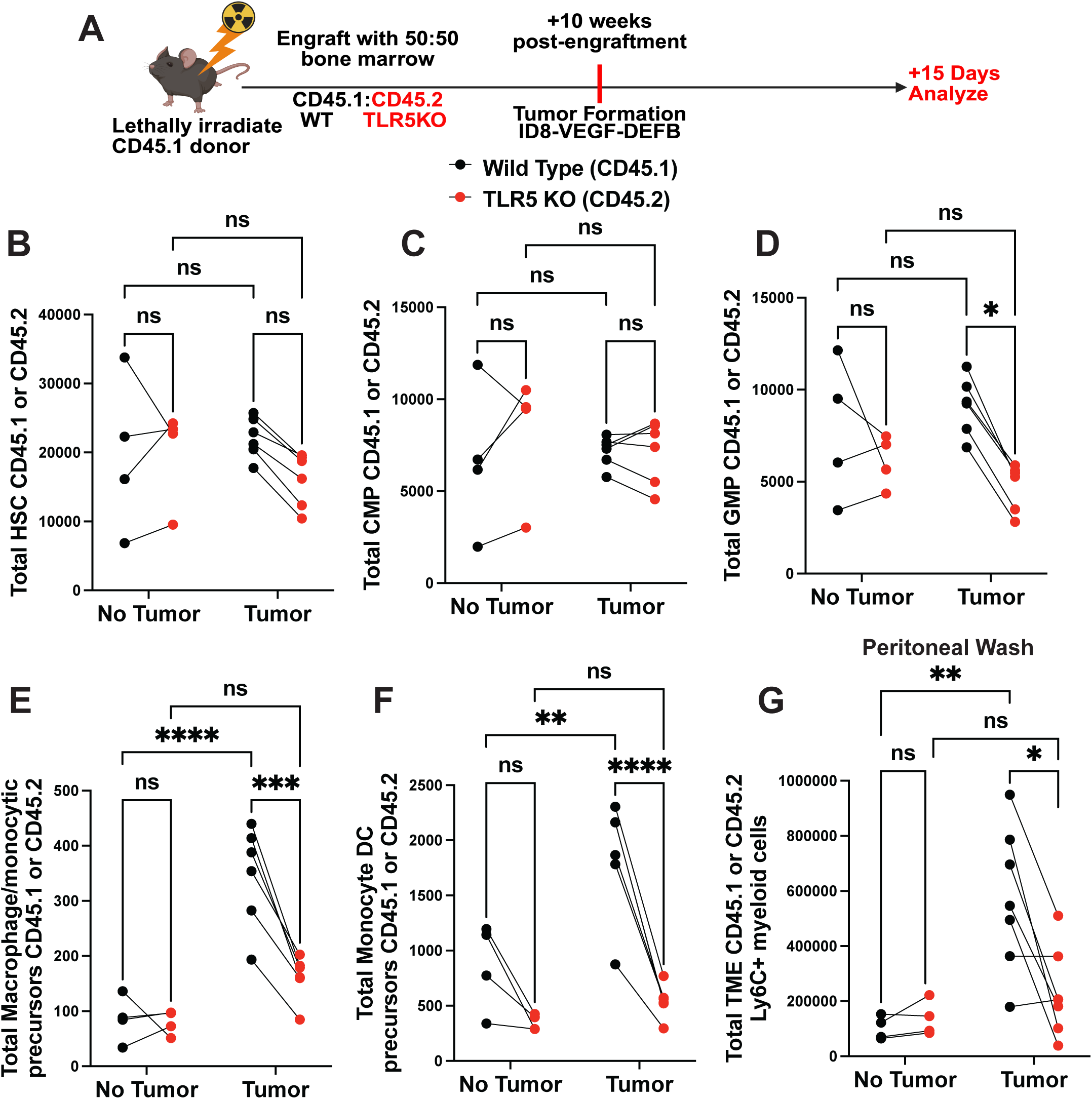
TLR5 signaling intrinsically drives myeloid progenitor differentiation and Ly6C⁺ myeloid accumulation in the ovarian tumor microenvironment. **(A)** Schematic of competitive bone marrow chimera experiment. WT CD45.1 recipient mice were lethally irradiated and reconstituted with a 1:1 mixture of wild-type (WT, CD45.1⁺) and TLR5 knockout (TLR5 KO, CD45.2⁺) donor bone marrow. After 10 weeks of engraftment, mice were either left untreated or implanted intraperitoneally with ID8-Defb29/VEGF-A ovarian tumors. Tissues were analyzed 15 days post-tumor implantation. **(B–F)** Quantification of hematopoietic stem and myeloid progenitor populations in the bone marrow of tumor-bearing and non-tumor-bearing mice. Each dot represents one mouse; lines connect WT and KO values from the same animal. **(B)** Hematopoietic stem cells (HSCs; Lin⁻ Sca-1⁺ c-Kit⁺) showed no significant differences between WT and TLR5 KO compartments regardless of tumor status. **(C)** Common monocyte progenitors (CMPs; Lin⁻ Sca-1⁻ c-Kit⁺ CD34⁺ CD16/32^int) were also unchanged between genotypes across conditions. **(D)** Granulocyte-monocyte progenitors (GMPs; Lin⁻ Sca-1⁻ c-Kit⁺ CD34⁺ CD16/32^hi) were significantly increased in WT versus TLR5 KO compartments in tumor-bearing mice, indicating that tumor-induced GMP expansion requires TLR5 expression. **(E)** Macrophage/monocytic precursors (CD11b⁺ Ly6C⁺ CCR2⁺ F4/80⁻) were enriched in WT cells compared to TLR5 KO under tumor conditions. **(F)** Monocyte dendritic cell precursors (CD11b⁺ Ly6C⁺ CD11c⁺ F4/80⁻) were similarly increased in WT cells in tumor-bearing mice. **(G)** Total Ly6C⁺ myeloid cells (CD11b⁺ Ly6C⁺) within the tumor microenvironment were significantly higher in the WT compartment compared to TLR5 KO, linking bone marrow-intrinsic TLR5 signaling to myeloid cell accumulation in the tumor. Data represent paired measurements from chimeric mice. Bars indicate mean ± SEM. Statistics: paired two-tailed t-tests. *p < 0.05, **p < 0.01, ***p < 0.001, ****p < 0.0001; ns = not significant.

In the bone marrow, total numbers of hematopoietic stem cells (HSCs) and common monocyte progenitors (CMPs) did not differ between WT and TLR5 KO compartments under either tumor-free or tumor-bearing conditions (**Figure 4B–C**). However, granulocyte-monocyte progenitors (GMPs) were significantly expanded in the WT compartment of tumor-bearing mice compared to their TLR5-deficient counterparts (**Figure 4D)**, indicating a TLR5-intrinsic requirement for expansion of GMP in response to tumor challenge. This corresponded to an expansion of cells further downstream of the monocytic lineage. In tumor-bearing mice, WT macrophage/monocytic progenitors and monocyte-dendritic cell precursors were increased compared to those derived from TLR5 KO progenitors (**Figure 4E–F**). Importantly, these differences were not observed in non-tumor-bearing controls or in other non-monocyte progenitor populations such as common lymphocyte progenitors (**Supplemental Figure 3C**) or in megakaryocyte-erythroid progenitors (**Supplemental Figure 3D**), suggesting that the TLR5-dependent effect on bone marrow progenitors was specific to monocyte progenitor cells and was dependent upon the presence of a tumor.

To determine whether changes within the bone marrow corresponded to differences in monocyte populations within the TME, we assessed Ly6C+ myeloid cell populations in the TME of chimeric mice. We observed a significant expansion in WT Ly6C+ myeloid cells compared to those from TLR5 KO cells in tumor-bearing animals (**Figure 4G**). Interestingly we did not observe a difference in tumor-associated granulocytes between TLR5 KO and WT cells, suggesting that TLR5 signaling is promoting the expansion of monocytic progenitor cells, as opposed to a global expansion in all myeloid lineages or myeloid-derived suppressor cells. Overall, these data suggest that in tumor-bearing mice, TLR5 signaling promotes the expansion of GMPs and monocytic progenitors in the bone marrow which leads to an accumulation of monocytes and macrophages within the ovarian tumor microenvironment

### TLR5 Signaling Influences Hematopoietic Colony Formation in Response to Stimulation

Given that our previous findings demonstrated a role for TLR5 signaling in the expansion of myeloid progenitor cells, we next sought to determine whether direct TLR5 stimulation could influence the differentiation potential of hematopoietic progenitors. To test this, we performed methylcellulose colony-forming unit (CFU) assays using bone marrow isolated from WT and TLR5 KO non-tumor-bearing mice with or without bacterial flagellin or LPS (**Figure 5A**). LPS was included as a positive control based on prior studies demonstrating that TLR4 signaling drives the expansion of myeloid progenitors under inflammatory conditions(41).

**Figure 5.**
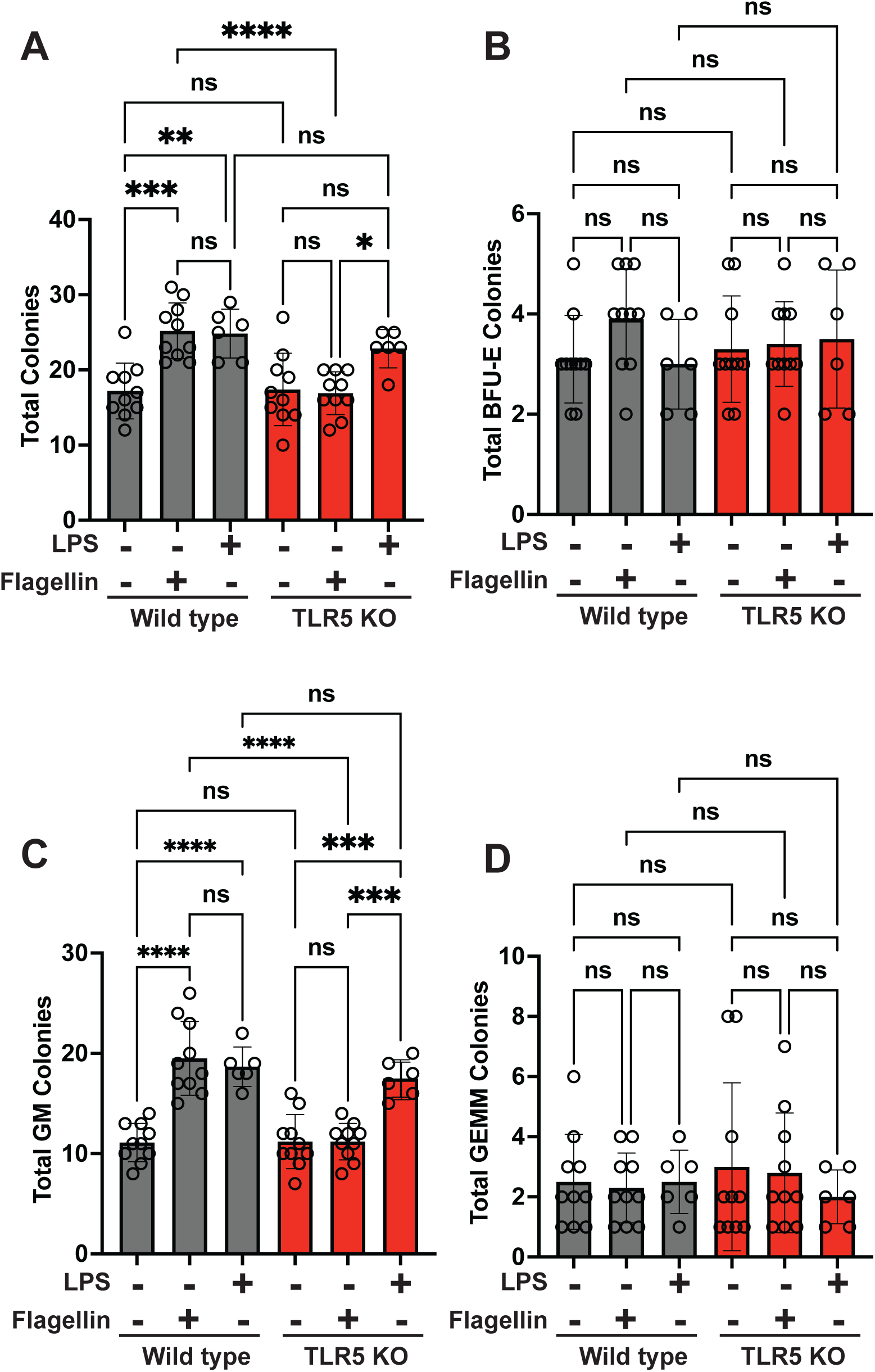
TLR5 signaling promotes myeloid-biased hematopoietic colony formation in response to bacterial ligands. (A–D) Quantification of colony-forming units (CFUs) generated from wild-type (WT, gray bars) and TLR5 knockout (TLR5 KO, red bars) mice in response to ultra-purified flagellin (TLR5 ligand), lipopolysaccharide (LPS, TLR4 ligand), or no stimulation. Whole bone marrow cells were isolated from non-tumor-bearing mice and plated into MethoCult M3434 methylcellulose medium, which supports multilineage hematopoiesis. Cultures were supplemented with 100 ng/mL flagellin (FLA-ST), 100 ng/mL LPS, or vehicle, and incubated for 11 days prior to manual colony quantification by morphology. **(A)** Total colony counts. Flagellin significantly increased total colony output in WT but not TLR5 KO cultures. LPS enhanced colony formation across both genotypes. **(B)** Burst-forming unit-erythroid (BFU-E) colony numbers were unchanged across all conditions. **(C)** Granulocyte-macrophage (GM) colonies were significantly elevated in WT cultures following either flagellin or LPS treatment, whereas TLR5 KO cells responded only to LPS. **(D)** Granulocyte, erythroid, monocyte, megakaryocyte (GEMM) colony numbers were not significantly altered by stimulation or genotype. Statistical comparisons were made using one-way ANOVA with multiple comparisons; *p < 0.05, **p < 0.01, ***p < 0.001, ****p < 0.0001; ns = not significant. Bars represent mean ± SEM; each point represents an individual mouse.

Stimulation of WT bone marrow with flagellin significantly increased the total number of colonies compared to unstimulated controls, whereas no such effect was observed in TLR5 KO cultures stimulated with flagellin (**Figure 5A**). On the other hand, LPS stimulation increased total colony formation in both WT and TLR5 KO cells, consistent with a TLR4-dependent mechanism. When evaluating colony subtypes, we observed no significant changes in burst-forming unit-erythroid (BFU-E) or granulocyte, erythrocyte, monocyte, megakaryocyte (GEMM) colonies across genotypes or conditions (**Figure 5B, 5D**). The most pronounced effect was observed in granulocyte-macrophage (GM) colonies. Both flagellin and LPS significantly increased GM colony formation in WT cells, while only LPS induced a similar effect in TLR5 KO cells (**Figure 5C**). This suggests that TLR5 signaling specifically promotes skewing of progenitors toward the GM lineage in response to flagellin. Although both flagellin and LPS can expand GMP-derived colonies *in vitro*, prior work has demonstrated that in the context of ovarian cancer progression, TLR4 signaling does not influence immune suppression and failure of immune therapy, whereas the effect is specific to TLR5 signaling(1). In support of these findings, previous studies have shown that TLR5 signaling can stimulate myeloid progenitors to mobilize in peripheral tissues such as the lung(42), and prostate(43). This suggests that TLR5 engagement is not only sufficient to promote myeloid bias *in vitro* but may also guide the recruitment and expansion of these cells *in vivo*.

## Discussion

In this study, we identify a novel role for TLR5 signaling in the expansion of monocyte progenitors, which leads to the accumulation of these cells within the ovarian TME. We first demonstrate that ovarian cancer induces gut barrier disruption, permitting the systemic translocation of TLR5 ligands into distal tissues such as the peritoneal cavity, blood, and bone marrow. These effects occur independently of host TLR5 expression and mirror those observed following DSS-induced colitis. Using a TLR5 reporter cell line, we quantified the presence of TLR5 ligands and confirmed elevated levels in tumor-bearing mice. Building on previous findings that TLR5 is primarily expressed on myeloid populations within the ovarian TME (2) (1), we show that pharmacologic blockade of TLR5 signaling significantly reduces the composition of tumor-infiltrating myeloid cells, culminating in a reduction of Ly6C+ monocytes and macrophages. In the bone marrow, TLR5 KO mice do not exhibit tumor-induced expansion of monocytic progenitors to the level of that observed in WT mice. Mixed bone marrow chimera experiments further revealed that TLR5 signaling leads to an expansion of granulocyte-monocyte progenitors and monocytes in tumor-bearing mice. Finally, stimulation of bone marrow progenitors with ultra-purified flagellin promotes the formation of myeloid-biased colonies in WT but not TLR5 KO bone marrow progenitors, further implicating that direct TLR5 signaling is driving the expansion or differentiation of monocytes. Together, these data support a model in which ovarian cancer–associated gut leakage and systemic dissemination of microbial ligands promote chronic TLR5 signaling, which acts at both the level of the bone marrow and the TME(1) to expand and polarize myeloid populations that are associated with immune suppression and poor outcomes across multiple cancer types(37,38).

These findings build on existing literature describing the immunosuppressive architecture of the ovarian TME, where myeloid-derived cells, particularly immature and inflammatory monocytes and tumor-associated macrophages, contribute to poor prognosis and immune evasion. Prior work has shown that TLR5 is predominantly expressed on CD11b+ and CD11c+ myeloid subsets within the ovarian TME, with minimal expression on T cells, suggesting that its effects are largely myeloid-intrinsic(1,2). In our study, the accumulation of Ly6C+CCR2+ monocytes following tumor growth was reduced in contexts where TLR5 signaling was limited, pointing to a role for TLR5 signaling in shaping both the recruitment and differentiation of myeloid populations. The presence of TLR5 ligands in the bone marrow further supports a systemic axis by which gut leakage promotes early myeloid skewing prior to infiltration into the tumor. This aligns with other studies, including in pulmonary models, showing that TLR5 activation promotes the migration and differentiation of progenitor cells into macrophages(42).

Beyond expanding myeloid populations, our findings suggest that TLR5 signaling may influence their functional polarization within the TME. Prior work by McGinty et al. demonstrated that TLR5-expressing dendritic cells in ovarian tumors exhibit reduced cross-presentation capacity and increased expression of inhibitory ligands, contributing to an immunosuppressive milieu(1). This adds to a growing body of evidence implicating pattern recognition receptor signaling in the modulation of myeloid cell fate. For instance, TLR7 activation has been shown to polarize myeloid cells toward immunoregulatory phenotypes in certain cancer settings(44). Similarly, TLR5 signaling has been reported to drive the polarization of tumor-associated macrophages in pancreatic cancer(43), further supporting its broader immunomodulatory capacity. Together, these findings position TLR5 as a dual regulator of both myeloid expansion and functional skewing within tumors, which form the basis of our future studies.

Importantly, while both TLR5 and TLR4 ligands flagellin and LPS, respectively, can expand granulocyte-monocyte progenitors *in vitro*, our data and previous work demonstrate that only TLR5 signaling is associated with immune suppression in the context of ovarian cancer(1). In methylcellulose assays, LPS stimulated colony formation in both WT and TLR5-deficient progenitors, whereas flagellin-induced colony formation was strictly TLR5-dependent. However, for ovarian cancer, only TLR5 expression corresponded with the promotion of an immune suppressive TME and failure of immune therapies(1). These findings suggest that the contribution of TLR5 to ovarian cancer is not merely due to its ability to expand myeloid progenitors but also reflects its role in programming the immune-suppressive characteristics of these cells within the TME. These data suggest that TLR5, but not TLR4, plays a unique and detrimental role in shaping the myeloid compartment during tumor progression for ovarian cancer. Differences in receptor expression patterns on affected cell populations, ligand bioavailability, and potential synergistic signaling partners may underlie this divergence. While tools to directly compare TLR4 and TLR5 ligand dissemination and receptor-specific signaling *in vivo* remain limited, our findings emphasize that TLR5 has a distinct, tumor-promoting function in ovarian cancer that is not recapitulated by TLR4 signaling.

Together, these findings position TLR5 as a key regulator of tumor-associated myelopoiesis, operating at the intersection of barrier disruption, microbial sensing, and immune suppression. By linking gut leakage to altered hematopoietic output in the bone marrow, our work extends prior understanding of how systemic inflammation contributes to the evolution of the ovarian tumor microenvironment. While pattern recognition receptor signaling has long been implicated in cancer immunity, our data underscore the specificity of TLR5 as a driver of myeloid and monocyte expansion and accumulation within the TME, with limited redundancy from other similar pathways such as TLR4. Future studies leveraging lineage-specific TLR5 deletion models and paired transcriptomic profiling of myeloid subsets will be essential to further dissect the signaling events that govern TLR5-mediated myeloid bias and immune suppression in ovarian cancer.

## Supporting information

Kolli et al Supplemental

