## Supplementary material for "Ovarian Cancer Drives TLR5-Dependent Expansion of Myeloid Progenitors Through Systemic Dissemination of Ligands": Kolli et al Supplemental

### Supplementary Data

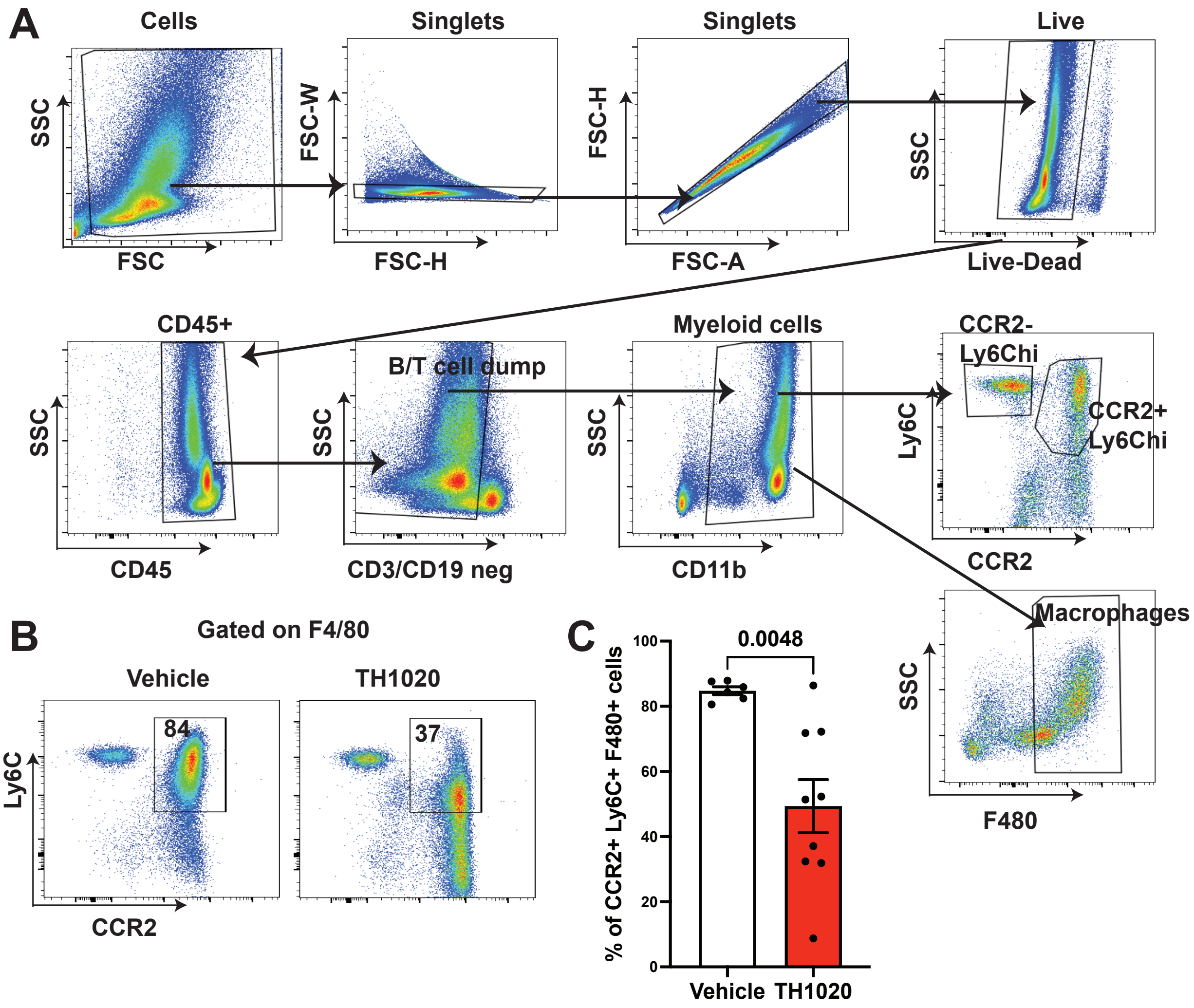


**Supplemental Figure 1: Gating strategy and flow cytometric analysis of CCR2$^+$ Ly6C$^+$ monocytic populations following TLR5 blockade in tumor-bearing mice. (A)** Representative gating strategy for flow cytometric identification of tumor-infiltrating CCR2$^+$ Ly6C$^+$ F4/80$^-$ myeloid cells. Single-cell suspensions were prepared from ascites of ID8-Defb29/VEGF-A tumor-bearing mice treated with either vehicle or the TLR5-neutralizing antibody TH1020. Live, singlet cells were first gated using forward and side scatter properties (FSC/SSC) and doublets excluded by FSC-H versus FSC-A. Live cells were identified by exclusion of the viability dye and gated on CD45$^+$ leukocytes. CD3$^-$CD19$^-$ cells were selected to exclude lymphoid lineages, and CD11b$^+$ myeloid cells were further gated for Ly6C and CCR2 expression. Final gating was performed on F4/80$^-$ cells to identify non-macrophage monocytic cells within the CCR2$^+$ Ly6C$^+$ compartment. **(B)** Representative flow cytometry plots showing frequency of CCR2$^+$ Ly6C$^+$ F4/80$^-$ cells from vehicle- and TH1020-treated mice. The percentage of gated events within the CCR2$^+$ Ly6C$^+$ population is indicated in the top left quadrant. **(C)** Quantification of CCR2$^+$ Ly6C$^+$ F4/80$^-$ myeloid cells as a percentage of total CD11b$^+$ myeloid cells in the tumor ascites. Treatment with TH1020 significantly reduced the proportion of these cells compared to vehicle-treated controls. Each dot represents an individual mouse. Bars indicate mean ± SEM. Statistical significance was assessed using an unpaired two-tailed t-test; $p = 0.0048$.

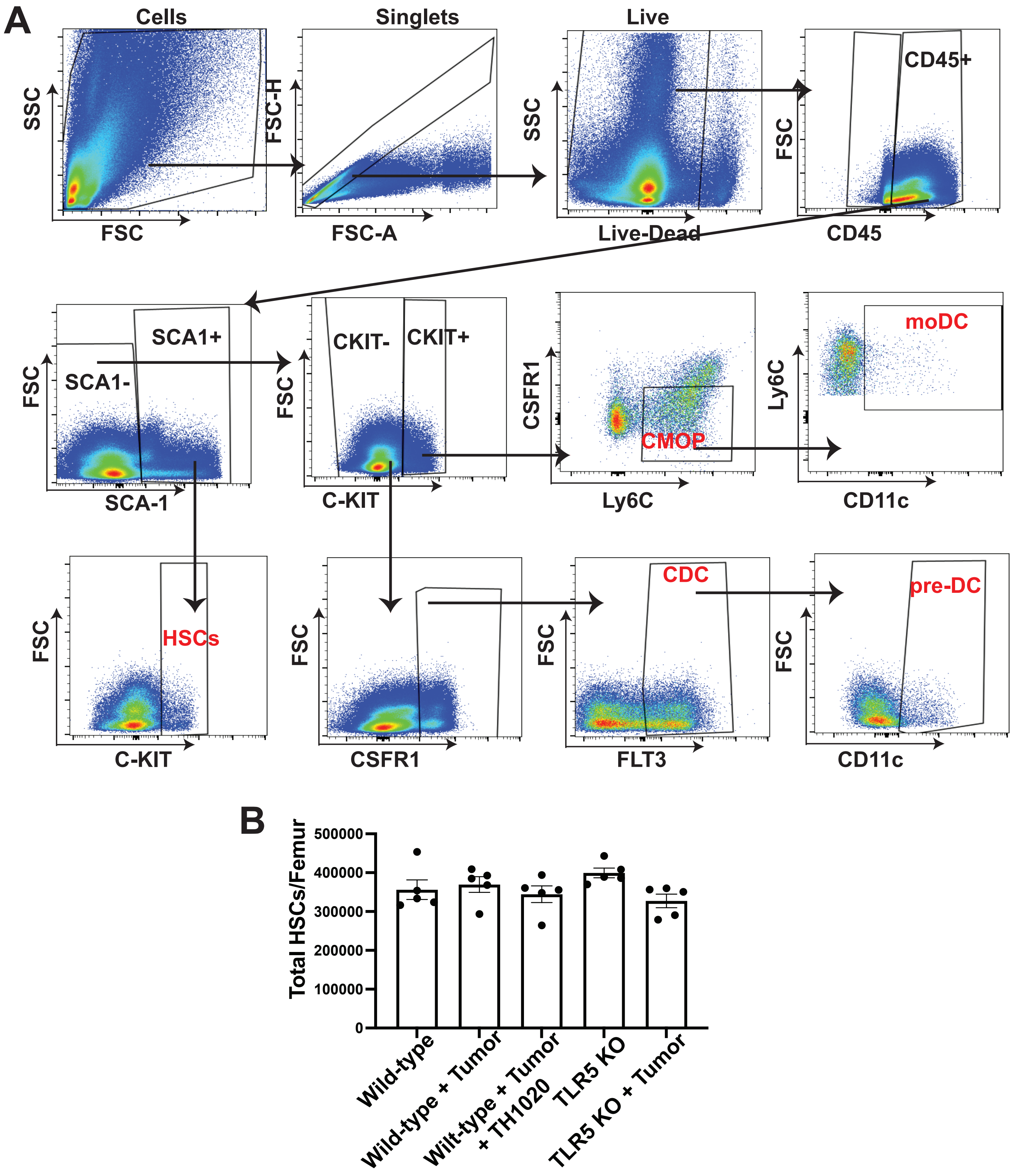
A
Cells
Singlets
Live
SSC
FSC
FSC-H
FSC-A
SSC
Live-Dead
FSC
CD45
CD45+
FSC
SCA-1
SCA1+
SCA1-
FSC
C-KIT
CKIT-
CKIT+
CSFR1
Ly6C
CMOP
Ly6C
CD11c
moDC
FSC
C-KIT
HSCs
FSC
CSFR1
FSC
FLT3
CDC
FSC
CD11c
pre-DC
B
Total HSCs/Femur
500000
400000
300000
200000
100000
0
Wild-type
Wild-type + Tumor
Wilt-type + Tumor + TH1020
TLR5 KO
TLR5 KO + Tumor

**Supplemental Figure 2: Gating strategy and quantification of hematopoietic stem cells (HSCs) in the bone marrow. (A)** Representative flow cytometry gating strategy for identification of hematopoietic stem and progenitor cell subsets from mouse femoral bone marrow. Single-cell suspensions were first gated to exclude debris and doublets using forward scatter area (FSC-A), height (FSC-H), and side scatter (SSC) parameters. Live cells were selected based on viability dye exclusion and further gated on $CD45^+$ hematopoietic cells. Lineage-negative cells ($CD3^-/CD19^-$) were used to isolate progenitor populations. HSCs were identified as $Lin^-$ $CD45^+$ $SCA\text{-}1^+$ $C\text{-}KIT^+$ cells. Common monocyte progenitors (CMOPs) were gated as $CSFR1^+$ $Ly6C^+$ cells, while monocyte-derived dendritic cell precursors (moDCs) were defined as $Ly6C^+$ $CD11c^+$. Classical dendritic cell precursors (CDCs) were defined as $CSFR1^+$ $FLT3^+$ cells, and pre-classical dendritic cells (pre-DCs) were gated as $FLT3^+$ $CD11c^+$ cells. **(B)** Quantification of total HSCs per femur across five experimental groups: wild-type (WT), WT + tumor, WT + tumor + TH1020, TLR5 KO, and TLR5 KO + tumor. No significant differences in HSC number were observed across any condition. Each point represents one mouse; bars indicate mean ± SEM. Statistical analysis was performed using one-way ANOVA.

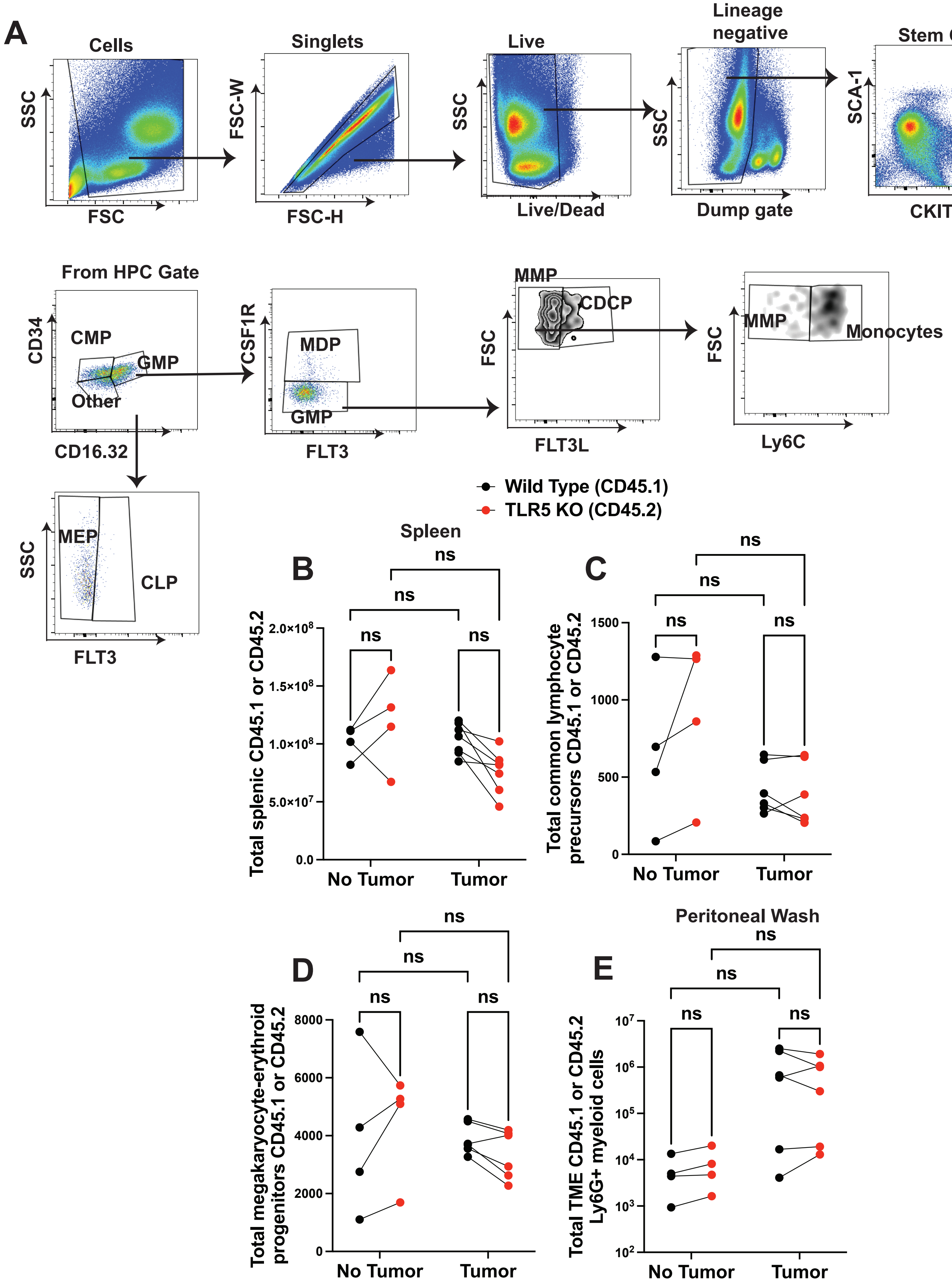
A
Cells
SSC
FSC
Singlets
FSC-W
FSC-H
Live
SSC
Live/Dead
Lineage negative
SSC
Dump gate
Stem Cells
SCA-1
CKIT
LSK
HPC
From HPC Gate
CD34
CD16.32
CMP
GMP
Other
CSF1R
FLT3
MDP
GMP
MMP
CDCP
FSC
FLT3L
MMP
Monocytes
FSC
Ly6C
MEP
CLP
SSC
FLT3
Wild Type (CD45.1)
TLR5 KO (CD45.2)
Spleen
B
Total splenic CD45.1 or CD45.2
No Tumor
Tumor
ns
C
Total common lymphocyte precursors CD45.1 or CD45.2
D
Total megakaryocyte-erythroid progenitors CD45.1 or CD45.2
Peritoneal Wash
E
Total TME CD45.1 or CD45.2 Ly6G+ myeloid cells

**Supplemental Figure 3: Gating strategy and systemic assessment of bone marrow chimera engraftment and immune cell subsets. (A)** Representative flow cytometry gating strategy for analysis of hematopoietic progenitors and myeloid subsets in competitive bone marrow chimeric mice. Single-cell suspensions were sequentially gated to exclude debris (FSC vs. SSC), doublets (FSC-H vs. FSC-W), and dead cells (viability dye). Hematopoietic stem and progenitor cells were gated as lineage-negative (dump gate: CD3, CD19, NK1.1, Ter119, Gr-1) SCA-1$^+$ C-KIT$^+$ (LSK) or SCA-1$^+$ C-KIT$^-$ cells. From the HPC gate, CMPs and GMPs were identified by CD34 and CD16.32 expression. Myeloid dendritic progenitors (MDP), granulocyte-monocyte progenitors (GMP), and common dendritic progenitors (CDCP) were resolved based on CSF1R, FLT3, and FLT3L expression. Monocyte populations were distinguished from MMPs based on Ly6C expression. Additional erythroid and lymphoid progenitors, including MEP and CLP, were gated based on FLT3 and SSC profiles.(B–E) Quantification of cell populations in wild-type (CD45.1) and TLR5 knockout (CD45.2) donor-derived cells from mixed bone marrow chimeras. Each line connects matched CD45.1 and CD45.2 values from the same mouse. **(B)** Total splenic CD45.1$^+$ and CD45.2$^+$ cells show no difference in engraftment in either tumor-bearing or tumor-free mice. **(C)** Total common lymphoid precursors (CLPs) from bone marrow demonstrate no significant difference between WT and TLR5 KO origin in either condition. **(D)** Megakaryocyte-erythroid progenitor (MEP) frequencies show no difference across genotype or tumor status. **(E)** Quantification of total Ly6G$^+$ myeloid cells in the tumor microenvironment (TME), stratified by CD45.1 and CD45.2 origin, reveals no significant differences, indicating equal recruitment and myeloid accumulation across genotypes. Data are pooled from ≥2 independent experiments. Each point represents an individual animal. Statistical analysis was performed using paired t-tests or nonparametric equivalents. ns = not significant.
